# Predictions and errors are distinctly represented across V1 layers

**DOI:** 10.1101/2023.07.11.548408

**Authors:** Emily R Thomas, Joost Haarsma, Jessica Nicholson, Daniel Yon, Peter Kok, Clare Press

## Abstract

‘Predictive processing’ frameworks of cortical functioning propose that neural populations in different cortical layers serve distinct roles in representing the world. There are distinct testable theories within this framework that we examined with a 7T fMRI study, where we contrasted responses in primary visual cortex (V1) to expected (75% likely) and unexpected (25%) Gabor orientations. Multivariate decoding analyses revealed an interaction between expectation and layer, such that expected events could be decoded with comparable accuracy across layers, while unexpected events could only be decoded in superficial laminae. These results are in line with predictive processing accounts where expected virtual input is injected into deep layers, while superficial layers process the ‘error’ with respect to expected signals. While this account of cortical processing has been popular for decades, such distinctions have not previously been demonstrated in the human sensory brain. We discuss how both prediction and error processes may operate together to shape our unitary perceptual experiences.

## Introduction

Popular accounts of mind and brain propose that the brain continuously forms predictions about future sensory inputs, and combines predictions with inputs to determine what we perceive^1–5^ (see also^6^). Such processes likely rely upon the hierarchical organization of the cortex, whereby feedforward connections relay information from superficial layers of lower cortical regions to the middle layers of higher areas, while feedback connections relay information from higher-level deep layers to agranular (superficial and deep) lower-level layers^7–10^. The precise functional roles of these ascending and descending connections across the hierarchy are, however, topics of debate.

‘Predictive processing’ schemes propose that feedback connections mediate contextual guidance of sensory processing by relaying predictions about the current state of the world, while feedforward connections convey sensory input^11,12^. At each level of the cortical hierarchy, predictions are compared to the input to compute the ‘prediction error’, which is transmitted up the hierarchy to update higher-level predictions until errors are reconciled^11,13–16^. A key feature of this account is that predictions and errors are represented by distinct neural populations residing in different cortical layers. For example, in primary visual cortex (V1), predictions conveyed via feedback projections have been proposed to influence representations in hypothesis units that reside in deep layers while error signals may be computed in superficial layers (Fig. 1A).

**Figure 1.**
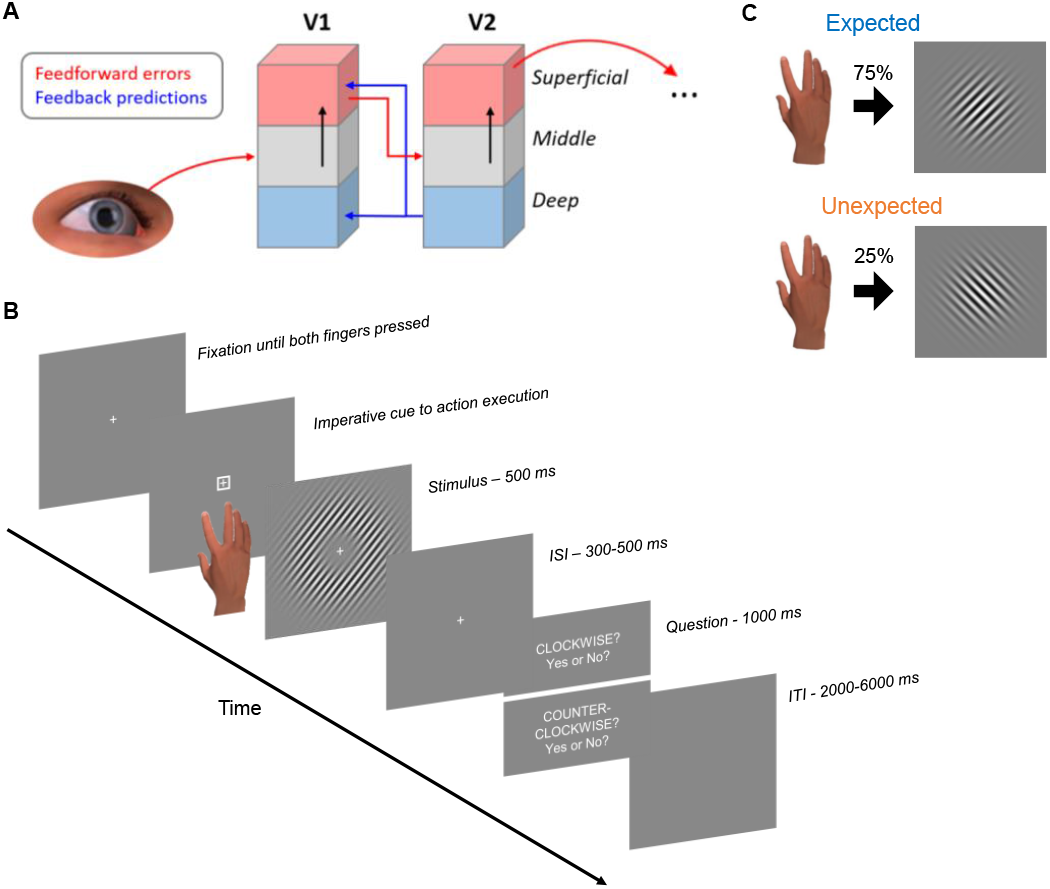
(A) Schematic representation of proposed extrinsic feedforward (red) and feedback (blue) connections across layers in early visual areas. (B) Experimental paradigm. A centrally presented visual cue instructed participants to abduct either their index or little finger. Each finger abduction predicted an oriented Gabor and participants were required to respond (yes/no) to whether the stimulus was clockwise (CW) or counter-clockwise (CCW) oriented relative to the vertical. In the training phase, actions perfectly predicted the stimulus orientation (100% contingency). (C) In the scanning session 24hrs later, participants completed the same task, but the action-outcome relationship was degraded to 75% to produce unexpected (25%) as well as expected (75%) events.

Despite the framework’s popularity, there is little understanding of functional distinctions, particularly because, to our knowledge, unexpected sensory events have not previously been presented in human laminar paradigms to contrast against expected events. This represented the aim of the present study. There are at least two mechanisms that could support the integration of our predictions with sensory input. First, a number of fMRI studies have observed improved sensory decoding of events expected on the basis of preceding cues, relative to unexpected events^17–20^, which may suggest that channels tuned to expected inputs are more responsive than channels tuned to the unexpected. Such a ‘global sharpening’ account could be supported if the precision weights are adjusted to increase the gain of expected channels, allowing them to respond more sensitively to ascending input^21^. Under this view, input channels may be more responsive to expected input, subsequently improving representation of the expected across the cortical column. If this were true, the unexpected would be represented poorly across layers as one effectively inhibits channels that report unexpected news from the sensory world. This account may explain biases to perceive what we expect^1–5^, if we relatively inhibit processing via other channels across layers.

An alternative proposes that future-based (e.g. cue-based) predictions can act like those about the present, such that they inject virtual input into hypothesis units in deep layers^22^. Notably, the predictive processing framework in general is concerned with ‘predicting’ (or representing) the present state of the world^21^, but in principle future-based and present-state predictions could operate similarly. Such proposals are in line with evidence from human ultra-high resolution 7T MRI studies demonstrating activity in deep cortical layers of V1 for visual events that are expected but never presented^22,23^(see also^24^). Without the need to invoke separable adjustments of precision weighting, like in the above global sharpening account, this ‘interaction’ account predicts a larger error signal in superficial layers for unexpected events – because the deep layers do not explain away such activity. Such holding of an error signal separate from the prediction, even in early sensory processing, may represent a mechanism enabling rapid model-updating^4^.

These predictions were tested in the present study, examining distinct processing across cortical layers of V1 using 7T fMRI. Participants were trained with perfect relationships between finger actions and visual Gabor orientations (e.g., index finger abduction = clockwise oriented Gabor; little finger abduction = counter-clockwise Gabor). They were presented at test (scanning phase) with degraded contingencies to measure neural responses to expected (in line with perfect contingency training phase; 75% of trials in the scanner) and unexpected (25%) events. Linear support vector machines (SVMs) were trained to discriminate Gabor orientations from a localizer and were tested on the main task^25^. The global sharpening account predicts superior decoding of expected relative to unexpected events across all cortical layers, whereas the interaction account predicts an interaction between expectation and layer.

## Results

Twenty-two participants (17 female, mean age = 26.09 years, SD = 3.41) were presented with perfect relationships between index and little finger actions and visual Gabor orientations during a training session. The next day, they completed the test session (in a 7T MRI scanner) where they were presented with degraded contingencies to measure neural responses to expected (in line with perfect contingency training phase; 75% of trials in the scanner) and unexpected (25%) events (see Fig. 1B-C). On half the trials they were asked to give a yes/no response to whether the stimulus was oriented CW and on the other half they were asked whether it was oriented CCW. This design orthogonalised the Gabor orientation presentation from the response. Expected and unexpected events were decoded from V1 activation patterns across layers using linear SVMs (Fig. 2).

**Figure 2.**
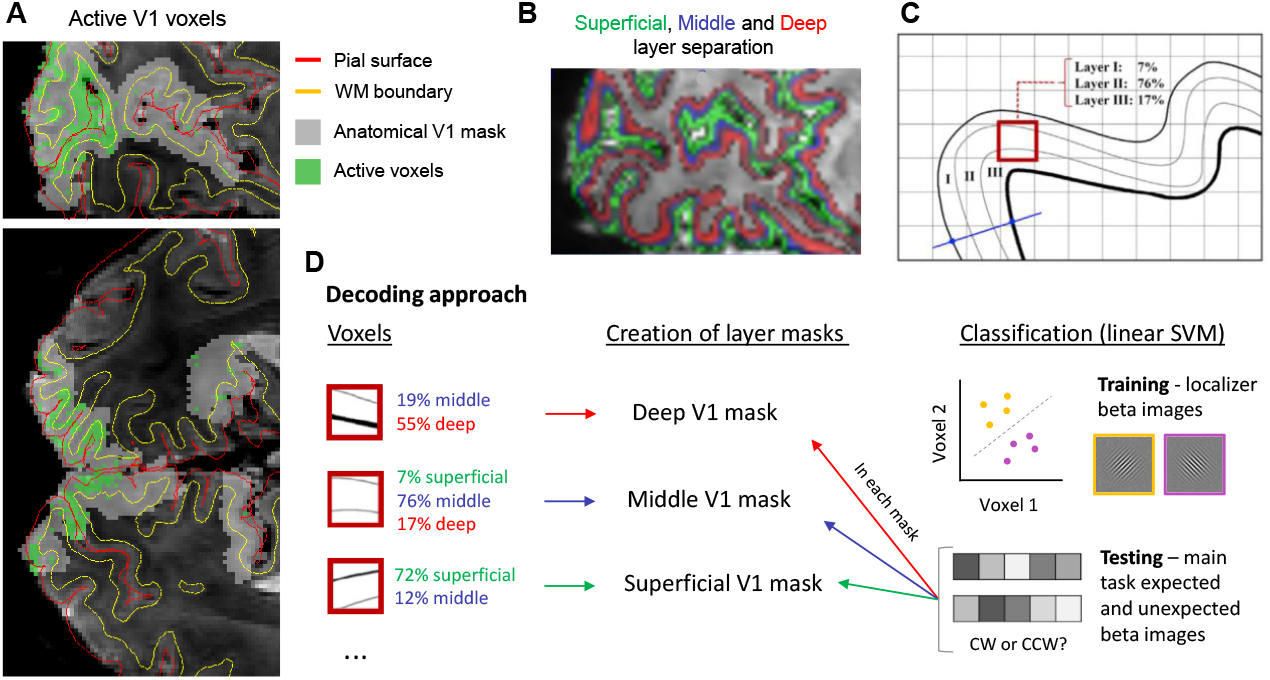
(A) Visualisation of the selected anatomical V1 ROI (light grey) on a mean functional image of an example participant. Overlaid red and yellow lines represent co-registered anatomical WM (yellow) and pial surface (red) boundaries to the mean functional image, showing voxels that were significantly active against baseline to the presented stimuli in the functional localiser task (green). (B) A mean functional image overlaid with distributions of voxels in superficial (green), middle (blue) and deep (red) layers of the cortex. (C) A schematic representing the level-set approach used to determine the volume distribution of a selected voxel (e.g. red square) over the superficial, middle, and deep cortical layers^22,26^. (D) A schematic of the decoding approach adopted here. Voxel proportions across the three layer bins in (C) were used to separate voxels according to the majority layer, and formed layer masks for V1. Linear classifiers (SVMs) were trained on CW and CCW stimuli from the localiser task and tested on Gabors from the main task. The procedure was repeated separately for expected and unexpected timecourses and in each V1 layer mask.

### Influence of expectations on behaviour

RT data were collected for responses to expected and unexpected stimuli in the test session and median RTs were calculated for correct trials separately for each condition and participant. Similarly, the proportion of correct responses was analysed for expected and unexpected conditions. RT analyses revealed no difference between expected (*M* = 586.78 ms, *SD* = 73.42) and unexpected (*M* = 589.98 ms, *SD* = 75.60) trials (*t*(19) = -.57, *p* = .58, *d* =-.13). Participants were however more accurate on expected (*M* = .97, *SD* = .03) than unexpected (*M* = .95, *SD* = .04) trials (*t*(19) = 2.67, *p* = .015, *d* = .60; see Fig. 3A).

**Figure 3.**
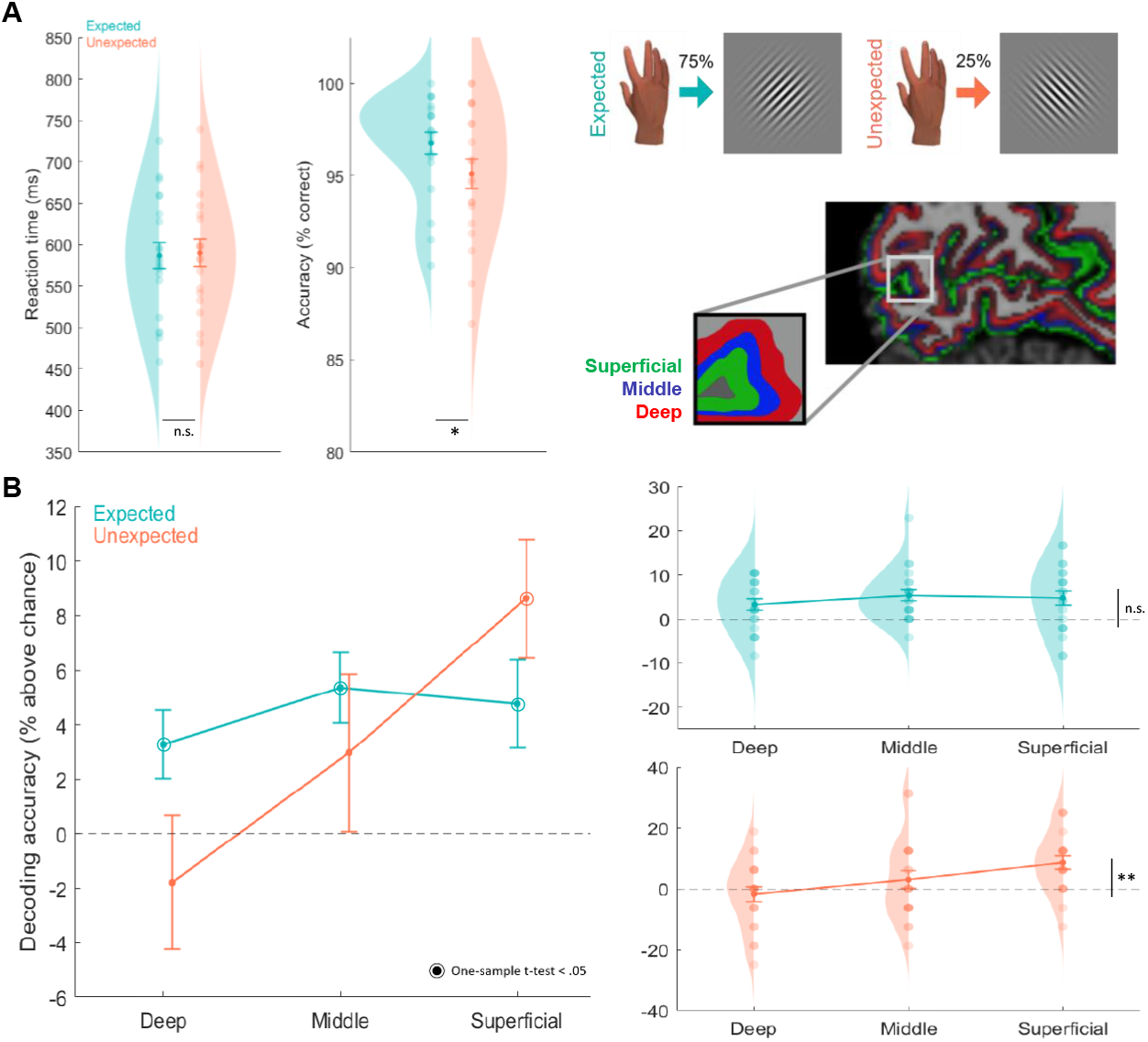
(A) Mean RTs and accuracy (± SEM) for expected and unexpected events, alongside probability density estimates and individual participant data points. There was no difference between conditions in RT, but participants were more accurate in expected than unexpected judgements (*p < 0.05). (B) The results of the decoding analysis across cortical layer bins, where mean (± SEM) decoding accuracy percentage above chance (50%) is plotted for expected and unexpected trials. On the left panel, circles around mean data points indicate that the decoding accuracy was significantly above chance (p < 0.05), which was the case across all layers for expected events but only in superficial layers for unexpected events. On the right panel, decoding accuracies are plotted alongside probability density estimates and individual participant data points. The linear trend across layers was significant for unexpected (**p < 0.001) but not expected events.

### Distinct cortical representations of predictions and errors

Using ultra-high field 7T fMRI (spatial resolution: 0.8-mm isotropic), we examined the brain activity patterns across cortical layers of V1 for expected and unexpected Gabor orientations. To determine layer-specific activity patterns, cortical V1 voxels were divided into 3 equivolume grey matter layer bins – superficial, middle and deep. The proportion of each voxel’s volume across the layers was used to create three cortical layer V1 masks for each participant that were used as layer ROIs for the following decoding analyses.

To investigate how expectations are represented across the cortical column in V1, linear SVMs were trained to discriminate Gabor orientations (CW or CCW) from a short localizer task that presented blocks of high-contrast flickering Gabors, and were tested on the main task (beta images from a first-level GLM). Given that expected events were presented with 75% likelihood, while unexpected events were presented with 25% likelihood, modelling all expected trials would render regressors that contained three times the data of unexpected regressors. We therefore modelled three expected regressors, all with an identical number of trials to unexpected regressors in the GLM, to reduce decoding biases across comparisons. The accuracy scores for each of the three expected decoding conditions were averaged for each participant, providing one accuracy score for ‘expected’ trials, and one for ‘unexpected’ trials, in each of the layer masks. These decoding accuracies were compared using a repeated measures ANOVA.

This analysis revealed a main effect of Layer (*F*(1.45,29.04) = 5.98, *p* = .012, *np*^2^ = .23, Greenhouse-Geisser corrected, ε = .73), no main effect of Expectation (*F*(1,20) = .39, *p* = .54, *np*^2^ = .019), but crucially, an interaction between Expectation and Layer (*F*(2,40) = 4.45, *p* =.018, *np*^2^ = .18; see Fig. 3B). Two control analyses were run at the voxel selection stage to balance the number of voxels in each layer mask, considering that there were more voxels in the superficial layers (due to known draining vein biases in gradient-echo EPI^27–31^). The effects remained in both a voxel-balanced control *t*-map analysis (this analysis selected an equal number of the most active voxels across the three layer bins; see Methods; Expectation x Layer: *F*(2,40) = 3.64, *p* = .035, *np*^2^ = .15; Layer: *F*(1.60,31.98) = 4.83, *p* = .021, *np*^2^ = .20, Huynh-Feldt corrected, ε = .80; Expectation: *F*(1,20) = 1.77, *p* = .20, *np*^2^ = .08), and a random-sample analysis (selecting an equal number of voxels across layers sampled randomly from each layer; Expectation x Layer: *F*(2,40) = 4.35, *p* = .019, *np*^2^ = .18; Layer: *F*(2,40) = 4.70, *p* = .015, *np*^2^ = .19; Expectation: *F*(1,20) = 2.07, *p* = .17, *np*^2^ = .09).

This interaction was generated via relatively consistent decoding across layers for expected events (*F*(2,40) = .82, *p* = .45, *np*^2^ = .04), while decoding performance differed for unexpected events (*F*(2,40) = 6.90, *p* = .003, *np*^2^ = .26) – increasing from deep to superficial layers (Linear trend: *F*(1,20) = 27.34, p < 0.001, *np*^2^ = .58). These effects are complemented by one-sample t-tests demonstrating that decoding of expected events was significantly different from chance across all layers (deep: *t*(20) = 2.62, *p* = .016; middle: *t*(20) = 4.13, *p* =.001; superficial: *t*(20) = 2.94, *p* = .008), while unexpected events could only be decoded in superficial layers (deep: *t*(20) = -.73, *p* = .47; middle: *t*(20) = 1.03, *p* = .31; superficial: *t*(20) = 3.97, *p* = .001). Additional post-hoc tests revealed no significant differences between expected and unexpected decoding within each layer (Deep: *t*(20) = 2.01, *p* = .058, *d* = .44; Middle: *t*(20) = .94, *p* = .36, *d* = .21; Superficial: *t*(20) = -1.41, *p* = .18, *d* = .31), though the numerical differences reveal superior representation of expected events in deep (Expected: *M* = 3.27, *SD* = 5.73, Unexpected: *M* = −1.76, *SD* = 11.21) and middle layers (Expected: *M* = 5.36, *SD* = 5.95, Unexpected: *M* = 2.98, *SD* = 13.20), flipping to superior representation of the unexpected in superficial layers (Expected: *M* = 4.76, *SD* = 7.43, Unexpected: *M* = 8.63, *SD* = 9.98).

Taken together, these post-hoc tests demonstrate that representation of unexpected events increases towards the superficial layers of V1, only becoming significantly decodable in these layers, while expected events are represented similarly across the cortical column.

## Discussion

The present study examined how unexpected visual events are represented across cortical layers, in comparison with expected events, in a high-resolution fMRI study. It found, in line with previous work^22^, that expected events were represented (decoded) equivalently across deep, middle and superficial bins, but more novelly, that unexpected events were represented with varying fidelity – such that they were poorly represented in deep and middle layers and could only be decoded above chance in superficial layers.

These findings are in line with predictive processing accounts whereby predictions inject virtual input into hypothesis units^21^. Feedback connections relay predictions about likely future inputs, while feedforward connections transmit the error between predictions and inputs up the hierarchy to update higher-level predictions until errors are reconciled^11,13,15,16^. Importantly, it is proposed that predictions and errors are represented distinctly across cortical laminae, such that predictions conveyed via feedback projections reside in deep layers while superficial laminae represent the error signal^13^. The interaction between expectation and layer established in this dataset is therefore as anticipated under this account, where superficial layers play a key role in the processing of the error signal. This finding is also in line with mouse data indicating superficial layer discrepancy signals with respect to other types of feedback^32,33^ (see also^34,35^). While this account of cortical processing has been popular for a couple of decades, such distinctions have not previously been demonstrated in human cortical processing.

Superficial layer responses observed here to unexpected events should be considered in light of previous lower resolution (3T) human fMRI work that demonstrates superior representation of unexpected visual events in visual regions relative to expected events^36–38^ (see also^39^). These studies directly contrast with previously mentioned studies that demonstrate superior decoding of expected events relative to the unexpected^17–19^. While it is currently unclear how to explain these differences^4^, they could be informed by the current findings demonstrating that unexpected representation differs across the column. The specific neural effect observed when examining activity across layers at lower resolutions may depend upon relative precision of expectations and inputs, leading to differential weighting of error signals^4^.

The findings are less consistent with accounts whereby the expected is represented with greater fidelity across the cortical column^21^, previously proposed to account for findings – including of ours – that the expected is represented with greater sensory fidelity than the unexpected^17–19^. This account suggested that precision weights of channels are adjusted according to future-based predictions, such that expected channels are released from inhibition relative to unexpected channels. Given the present findings, an alternative way to conceptualise enhanced decoding of expected events at lower-resolution fMRI^36–38^ is that predictions and inputs relate to the same visual event in expected conditions, but not unexpected conditions. When predictions and inputs reflect different events, unexpected events may be decoded poorly because they are only represented in the error units in superficial layers. However, such errors can be decoded distinctly in superficial layer activity when it is separated from deep (and middle) activity with high-resolution laminar fMRI.

Forging functional conclusions about the operation of mechanisms across cortical layers has become possible with high resolution MRI, but is of course also plagued by interpretational issues due to venous draining of blood towards the pial surface^27–31^. Specifically, gradient echo BOLD signal is known to exhibit strong contributions from large veins situated perpendicular to the cortical surface as venous blood is drained from lower to upper cortical layers^16,28,30^. Such venous issues render it likely, for instance, that neural effects at deeper cortical layers contribute to responses in superficial layers^40^. Regardless of these methodological challenges, it is essential that we attempt to overcome them if we are to test these hypotheses concerning error encoding. By comparing responses between stimuli that are identical, other than their expectedness due to a preceding cue (as in the present study), we can mitigate such contributions – because venous draining influences should be equivalent for both expected and unexpected events. While expected events could be decoded across all layers, unexpected events could only be significantly decoded in superficial layers, suggesting that influences from such draining effects were somewhat mitigated using this approach – since unexpected information should also be decodable in deep and middle layers with draining effects.

Such distinct representation of prediction and error may be an adaptive solution allowing predictions to shape perception to serve a number of functions. Some of us have recently discussed how predictions often need to exhibit quite distinct behavioural shaping of perception to serve the organism^4,41^. To overcome noise in sensory processing and generate broadly accurate experiences rapidly, we may bias perception towards what we expect^42,43^. However, larger error signals (that cannot have resulted from noise) may require high perceptual resources, to enable accurate perception and resultant model updating. If we hold the error signal separate from the prediction, even in early sensory processing, this may be one way to enable these large error signals to communicate deviation rapidly to systems mediating model-updating – such as the locus coeruleus^44^. Future work must establish how these error signals relate to perception and model-updating to truly test these accounts, and examine whether error signals in superficial layers are calculated in the first feedforward sweep^45^ or subsequent stimulus processing iterations.

It has been suggested in various theoretical accounts that symptoms of psychosis, like hallucinations and delusions, can be explained in terms of aberrant signalling of prediction error as well as overweighting of expectations^46–48^. As demonstrated in the present study, laminar fMRI is capable of distinguishing the representation of these signals across different cortical layers. Therefore, laminar fMRI would be well suited to test the theoretical predictions from predictive coding models of psychosis, as well as comparing these mechanisms in other clinical and neurological populations characterized by aberrant perceptual inference, like Parkinson’s disease^49^.

In conclusion, the present study provides evidence that expected and unexpected visual events are distinctly represented across the cortical column in V1, via a novel 7T fMRI design that presented unexpected visual events alongside expected counterparts. Expected events were represented similarly across layers but unexpected events were only represented well in superficial layers. These findings contribute to our understanding of how predictions can interact with sensory inputs to shape what we perceive and how we interact with the world.

## Methods & Analyses

### Participants

Twenty-two participants were recruited (17 female, mean age = 26.09 years, SD = 3.41) from Birkbeck, University of London and UCL, and paid a small honorarium for participation. All participants reported normal or corrected to normal vision and had no history of psychiatric or neurological illness. One participant’s data were excluded due to a technical error during acquisition, which meant that event onsets in one run could not be modelled. This resulted in a final sample of 21 participants. The experiment was approved by the UCL ethics committee.

### Stimuli

Sinusoidal grating stimuli were created using MATLAB and presented against a grey background using Cogent Graphics. During training, stimuli were presented on a 14” Dell Laptop via an LCD screen (resolution: 1280×1024; refresh rate: 60 Hz) at a viewing distance of 45 cm, and during scanning on an LCD monitor (resolution: 1280×1024; refresh rate: 60 Hz) through a mirror at a viewing distance of 91 cm. In both sessions, stimuli were viewed at 15 degrees of visual angle. A Gaussian filter enveloped the grating stimuli to create Gabor patches of 80% Michelson contrast, at 1.5 cycles per degree, and with random spatial phase. The Gabor stimuli were presented in an annulus around a fixation cross in the middle of the screen (see Fig. 1B). Two stimulus orientations were generated to appear in CW (45°) and CCW (135°) orientations (relative to the hypothetical vertical mid-point e.g., 90°).

### Procedure

#### Main task

Participants completed two sessions. First, they completed a training session in which finger abductions perfectly predicted visually presented Gabor orientations. The following day, they completed the same task in the MRI scanner but the action-outcome relationship was degraded to 75% validity to allow for presentation of unexpected (25%) as well as expected events.

Participants completed the training session on Gorilla (www.gorilla.sc) for online experiments, taking part on either a laptop or desktop computer no more than 24 hrs before the scanning session. Instruction at the beginning of the experiment requested participants to set screen brightness to the maximum level to reduce variability in viewing conditions. Each trial started with a white fixation cross. Participants were instructed to depress the ‘c’ and ‘m’ computer keys with their right index and little fingers, respectively, until an imperative cue (e.g., square or triangle overlaid around the fixation cross) indicated which finger to abduct. After the appropriate action was executed, the imperative cue was replaced with an oriented Gabor for 500 ms, resulting in apparent synchrony of stimulus onset with action execution. A variable 300 – 500 ms delay followed stimulus offset and preceded a response screen which asked about Gabor orientation. On half the trials they were asked to give a yes/no response to whether the stimulus was oriented CW and on the other half they were asked whether it was oriented CCW. This design orthogonalised the Gabor from the response. Participants were required to respond to the question screen within 1500 ms and the next trial started after a variable ITI of 2000-3000 ms. Responses were made using the left thumb on the ‘a’ and ‘z’ keys for ‘yes’ or ‘no’, respectively. The response question alternated every block. Participants completed the training task in ten runs of 36 trials each.

The following day, participants completed the test session at the Wellcome Centre for Human Neuroimaging, UCL. The test session task was largely similar to the training session except that participants’ abductions now predicted the stimulus orientation with 75% validity and they performed actions using MR-compatible button boxes instead of the keyboard. The right-hand button box was positioned orthogonally to the screen in the scanner, in line with the body midline. A short refresher of the training session was presented immediately before the scanning session, using the MR compatible button boxes outside of the scanner.

Responses were now required within 1000 ms of the question screen (to reduce scanning time), and the response question was randomly selected on each trial. The next trial started after a variable ITI of 2000-6000 ms. Participants completed the test session in four scanning runs that contained 96 trials each, and a 30 s break was presented mid-way through each run.

There were 384 main experimental trials in the scanning session, 360 online training trials and 192 refresher training trials. Participants completed 32 practice trials before proceeding to the main trials in the initial training session. Imperative cue order and trial order were randomised within blocks and the specific action-Gabor relationship was counterbalanced across participants. The imperative cue-action mapping was also counterbalanced and reversed halfway through each session (e.g., at the beginning of the sixth block in training, and beginning of the third block in scanning) to deconfound potential influences of cue-outcome learning and remove any correlation between the imperative action cues and actual or expected Gabor orientations across the experiment.

#### Localiser task

At the end of the main experiment, participants completed a functional localiser task in an additional scanning run. This task presented flickering Gabor stimuli at approximately 1.8 Hz along with a fixation cross. These Gabors were identical to those presented in the main experiment except that they were presented at 100% contrast, and in blocks of 14 s. Each block containing flickering Gabors was followed by a blank screen containing only the fixation cross for the same duration. In each stimulus block, Gabor orientation was either CW or CCW and the presentation order was pseudorandomised. The task required participants to respond by pressing any button when the central fixation cross changed colour from white to grey, ensuring that their fixation remained central. In total, 32 blocks of flickering Gabors were presented, 16 of each orientation.

### Behavioural analyses

RT data were collected for responses to expected and unexpected stimuli in the test session and median RTs were calculated for correct trials separately for each condition, for each participant. Similarly, the proportion of correct responses was analysed for expected and unexpected conditions for each participant. One participant was removed from the behavioural analysis due to missing almost half of the responses (44% of trials) and performing similarly to chance on the remainder (62% accuracy; note that this participant was maintained for the imaging analysis, but the significance patterns were identical if they were removed).

### Image Acquisition

Images were acquired using a 7T Magnetom MRI scanner (Siemens Healthcare GmbH, Erlangen, Germany) using a 32-channel head coil at the Wellcome Centre for Human Neuroimaging, UCL. Functional images were acquired using T2*-weighted 3D gradient-echo EPI sequence (3,552 ms volume acquisition time, TR = 74 ms, TE = 26.95 ms, 48 slices, 15° flip angle, voxel size: 0.8 × 0.8 × 0.8 mm, field of view: 192 × 192 × 39 mm). Structural images were acquired using a Magnetization Prepared Two Rapid Acquisition Gradient Echo (MP2RAGE) sequence (TR = 5,000 ms, TE = 2.60 ms, TI = 900 ms, 240 slices, voxel size 0.7 × 0.7 × 0.7 mm, 5° flip angle, field of view 208 × 208 × 156 mm).

### fMRI Data preprocessing

Preprocessing of the images was conducted in SPM12 and Freesurfer (http://surfer.nmr.mgh.harvard.edu/). Functional images were cropped to select only the occipital lobe, to account for distortions in the frontal lobes. These cropped functional images were spatially realigned to the mean image within runs, but also across runs. The temporal signal-to-noise ratio (tSNR, defined as mean signal/SD over time) was calculated before and after spatial realignment and was found to be significantly higher after (*M* =14.31, *SD* = 1.23) than before (*M* = 10.21, *SD* = 1.05) realignment (*t*(20) = −22.10, *p* < .001).

The realigned functional images were co-registered to the cortical surfaces estimated in participants’ MP2RAGE scans in several steps. First, boundaries between grey matter (GM), white matter (WM) and cerebrospinal fluid (CSF) were detected using Freesurfer on re-constructed structural scans (skull removed), and were manually corrected to remove any dura that was inaccurately classified as part of the GM surface. A rigid body boundary-based registration (BBR)^50^ was used to register GM boundaries to the mean functional image, and a further recursive boundary-based registration (RBR)^51^ applied the BBR recursively to portions of cortical mesh in 6 iterations.

### Cortical layer definition

The level set method was used to divide GM into three equivolume layers for cortical layer definition (for details see^26^). This method was used to separate five cortical bins (3 GM, WM and CSF) and determine three GM layers (deep, middle, and superficial) by calculating two intermediate surfaces between the WM and pial boundaries. In human V1, these three layer bins have been suggested to correspond to histological layers 1 to 3, layer 4, and layers 5 and 6, respectively^22^.

## Analyses

### Layer-specific ROI definition

Freesurfer was used to define V1 based on anatomical landmarks in the MP2RAGE scans. ROIs were restricted to voxels from the preprocessed functional localiser data that were most active during blocked presentation of the stimuli. This was achieved by modelling regressors for blocks of CW and CCW stimuli against baseline in a temporal GLM to identify voxels that expressed a significant response to these stimuli (*t* > 2.3, *p* < 0.05; *M* = 5420.57, *SD* = 2228.12 number of voxels). A V1 mask of active voxels was created this way for each participant.

Next, these active voxel masks were used to design a matrix of distributed voxels across each layer bin using the level-set definition described earlier^26^. These participant-specific design matrices specified the proportion of each active voxel across the 5 layer bins specified above (3GM, WM, CSF), where each voxel was binned into one of the three GM layer bins according to its majority proportion (Fig. 2D; see Supplementary Information for a complementary univariate analysis approach). For example, a voxel that was spatially located 7% in superficial, 76% in middle and 17% in deep layers would be labelled as a middle layer voxel and selected to contribute to the voxels in the middle layer mask. An arbitrary threshold was set such that the majority proportion for a voxel to be included in a layer mask was >0.4 (40%). This meant that any voxels with roughly equal proportion in each layer bin would not be selected. Importantly, the results did not change when this threshold was removed, since the majority of voxels’ ‘winning’ proportion was greater than 0.4. Using this approach, three layer masks were created from the active V1 voxels for each participant.

This method of defining layer specific ROIs yields V1 layer masks that differ in the number of voxels that contribute to each layer in each participant. Notably, there is a consistently greater number of voxels in superficial (*M* = 1613.95, *SD* = 678.17) than middle (*M* = 1172.48, *SD* = 530.94) and deep (*M* = 907.62, *SD* = 442.68) layer masks (one-way ANOVA: *F*(2,40) = 120.50, *p* < .001, *np*^2^ = .86). We therefore performed another analysis to control for these differences, considering that greater information contributing to the decoding signals in superficial layers relative to the other layers may confound our interpretations. Here, the steps are identical to above, except that an additional step was performed to equalise the number of voxels present in each layer ROI mask. Specifically, we defined the number of voxels to select in each mask as the maximum number common to all layers. For example, if the deep mask had the fewest and contained 831 voxels, 831 voxels would be selected across all layers. Next, we loaded in an orientation preference t-map from the GLM specified above, that contrasted CW and CCW regressors against each other, to select the (e.g., 831) most orientation-tuned voxels from each layer. These voxels were those that contributed to each layer mask, such that each layer mask contained an equivalent number of voxels in each layer. Another control version selected the (e.g. 831) voxels randomly from all the active voxels in each layer. Importantly, the results did not change across these selection methods, suggesting that differences in voxel numbers across layers should not alter interpretation (see Results).

### Decoding analysis

Multivariate decoding analyses were implemented using the TDT toolbox^52^ in MATLAB. We used a cross-classification approach whereby a linear SVM was trained to discriminate Gabor orientations (CW or CCW) presented during the localizer task. This independent dataset ensured that the trained classifier was not biased with any information about the predictability of stimuli. For this step, we reran the GLM that we used for ROI definition above, but instead specified the onsets for each block in the localizer task as separate regressors. Movement parameters were also modelled as nuisance regressors. This GLM resulted in 16 beta images for each orientation that were fed into the SVM for training.

Next, we specified the test data in our cross-classification decoding approach from our main experimental task data. Specifically, we reran and modified the GLM previously run on the main task data that included separate regressors for each condition type (expected, unexpected) and stimulus type (CW, CCW) in each experiment run, in two ways. First, considering that we only had 4 scanning runs, yet it is well established that decoding data is more reliable with increased number of samples, we modelled each condition according to the first and second halves of each run (since there was a 30s break in between continuous scanning; note also that all trial types were balanced within each run half). This resulted in 8 condition regressors for each scanning run (2x ExpCW1, ExpCCW1, UnexpCW1, UnexpCCW1). Second, we ensured that each modelled regressor would have equal weight in terms of the number of trials contributing to each image, considering known biases in decoding performance with unequal numbers of trials^53^. We therefore modelled expected conditions with the same number of trials as those that contribute to unexpected regressors, by randomly sampling from expected trials to form three different expected regressors (see Results). Again, movement parameters were modelled as nuisance regressors.

In total, this GLM resulted in 16 beta images (3x ExpCW, ExpCCW, 1x UnexpCW, UnexpCCW, twice in each scanning run). The beta images from this GLM were grouped according to our main experimental conditions such that we repeated the decoding procedure separately four times (Expected [x3], Unexpected) to determine whether stimulus orientation classification differed across each of these conditions. We tested each of these four decoding iterations separately, restricting the voxels to each of our three V1 layer masks. This procedure resulted in 12 testing iterations (4 conditions in each of 3 masks). Accuracy of the SVM was calculated as the proportion of correctly classified images across all decoding steps and was conducted separately for each participant. The accuracy scores for each of the three expected conditions were averaged for each participant, providing one accuracy score for ‘expected’ trials, and one for ‘unexpected’ trials, in each of the layer masks. These scores were then compared between expected and unexpected conditions, and across layers, to determine whether information about presented stimuli varied as a function of learned expectation across the cortical layer bins.

The results were then analysed with a 2×3 repeated measures ANOVA with the factors experimental condition (expected, unexpected) and cortical layer (deep, middle, superficial). Follow up tests examined differences across layers, separately for expected and unexpected events.

## Supporting information

Supplementary Methods and Results

## Acknowledgements

This work was supported by a Leverhulme Trust project grant (RPG-2016-105) and European Research Council (ERC) consolidator grant (101001592) under the European Union’s Horizon 2020 research and innovation programme, both awarded to CP. PK was supported by a Wellcome/Royal Society Sir Henry Dale Fellowship (218535/Z/19/Z) and an ERC Starting Grant (948548). ET and DY were supported by the Leverhulme Trust grant awarded to CP and JH by the ERC grant awarded to PK. The Wellcome Centre for Human Neuroimaging is supported by core funding from the Wellcome Trust (203147/Z/16/Z). We are grateful to Martina Callaghan for useful discussions.

## References

1. Bar, M. Visual objects in context. Nat. Rev. Neurosci. 5, 617–629 (2004).

2. Yuille, A. & Kersten, D. Vision as Bayesian inference: analysis by synthesis? Trends Cogn. Sci. 10, 301–308 (2006).

3. de Lange, F. P., Heilbron, M. & Kok, P. How do expectations shape perception? Trends Cogn. Sci. 22, 764–779 (2018).

4. Press, C., Kok, P. & Yon, D. The perceptual prediction paradox. Trends Cogn. Sci. 24, 13–24 (2020).

5. Den Ouden, H., Kok, P. & De Lange, F. How prediction errors shape perception, attention, and motivation. Front. Psychol. 3, (2012).

6. Kaiser, D., Quek, G. L., Cichy, R. M. & Peelen, M. V. Object vision in a structured world. Trends Cogn. Sci. 23, 672–685 (2019).

7. Felleman, D. J. & Van Essen, D. C. Distributed hierarchical processing in the primate cerebral cortex. Cereb. Cortex 1, 1–47 (1991).

8. Rockland, K. S. & Virga, A. Terminal arbors of individual “feedback” axons projecting from area V2 to V1 in the macaque monkey: A study using immunohistochemistry of anterogradely transported Phaseolus vulgaris-leucoagglutinin. J. Comp. Neurol. 285, 54–72 (1989).

9. van Kerkoerle, T., Self, M. W. & Roelfsema, P. R. Layer-specificity in the effects of attention and working memory on activity in primary visual cortex. Nat. Commun. 8, 1–14 (2017).

10. Yu, Y. et al. Layer-specific activation of sensory input and predictive feedback in the human primary somatosensory cortex. Sci. Adv. 5, eaav9053.

11. Friston, K. A theory of cortical responses. Phil Trans R Soc B 360, 815–836 (2005).

12. Keller, G. B. & Mrsic-Flogel, T. D. Predictive processing: A canonical cortical computation. Neuron 100, 424–435 (2018).

13. Bastos, A. M. et al. Canonical microcircuits for predictive coding. Neuron 76, 695–711 (2012).

14. Kanai, R., Komura, Y., Shipp, S. & Friston, K. Cerebral hierarchies: Predictive processing, precision and the pulvinar. Philos. Trans. R. Soc. B Biol. Sci. 370, (2015).

15. Rao, R. P. N. & Ballard, D. H. Predictive coding in the visual cortex: a functional interpretation of some extra-classical receptive-field effects. Nat. Neurosci. 2, 79–87 (1999).

16. Stephan, K. E. et al. Laminar fMRI and computational theories of brain function. NeuroImage 197, 699–706 (2019).

17. Kok, P., Jehee, J. F. M. & de Lange, F. P. Less is more: Expectation sharpens representations in the primary visual cortex. Neuron 75, 265–270 (2012).

18. Yon, D., Gilbert, S. J., De Lange, F. P. & Press, C. Action sharpens sensory representations of expected outcomes. Nat. Commun. 9, 4288 (2018).

19. Yon, D. et al. Stubborn predictions in primary visual cortex. J. Cogn. Neurosci. 35, 1133–1143 (2023).

20. Brandman, T. & Peelen, M. V. Objects sharpen visual scene representations: evidence from MEG decoding. bioRxiv (2023) doi:10.1101/2023.04.06.535903.

21. Friston, K. Does predictive coding have a future? Nat. Neurosci. 21, 1019–1021 (2018).

22. Aitken, F. et al. Prior expectations evoke stimulus-specific activity in the deep layers of the primary visual cortex. PLoS Biol. 18, e3001023 (2020).

23. Haarsma, J., Deveci, N., Corbin, N., Callaghan, M. F. & Kok, P. Perceptual expectations and false percepts generate stimulus-specific activity in distinct layers of the early visual cortex. bioRxiv (2022) doi:10.1101/2022.04.13.488155.

24. Muckli, L., De Martino, F., Ugurbil, K., Goebel, R., & Essa Yacoub. Contextual feedback to superficial layers of V1. Curr. Biol. 25, 2690–2695 (2015).

25. Peelen, M. & Downing, P. Testing cognitive theories using multivariate pattern analysis of neuroimaging data. Preprint at https://doi.org/10.31234/osf.io/rhzt9 (2022).

26. van Mourik, T., van der Eerden, J. P. J. M., Bazin, P.-L. & Norris, D. G. Laminar signal extraction over extended cortical areas by means of a spatial GLM. PLOS ONE 14, e0212493 (2019).

27. Lawrence, S. J. D., Formisano, E., Muckli, L. & De Lange, F. P. Laminar fMRI: Applications for cognitive neuroscience. NeuroImage 197, 785–791 (2019).

28. Duvernoy, H. M., Delon, S. & Vannson, J. L. Cortical blood vessels of the human brain. Brain Res. Bull. 7, 519–579 (1981).

29. Koopmans, P. J., Barth, M. & Norris, D. G. Layer-specific BOLD activation in human V1. Hum. Brain Mapp. 31, 1297–1304 (2010).

30. Markuerkiaga, I., Barth, M. & Norris, D. G. A cortical vascular model for examining the specificity of the laminar BOLD signal. NeuroImage 132, 491–498 (2016).

31. Heinzle, J., Koopmans, P. J., den Ouden, H. E. M., Raman, S. & Stephan, K. E. A hemodynamic model for layered BOLD signals. NeuroImage 125, 556–570 (2016).

32. Audette, N. J., Zhou, W., La Chioma, A. & Schneider, D. M. Precise movement-based predictions in the mouse auditory cortex. Curr. Biol. 32, 4925-4940.e6 (2022).

33. Jordan, R. & Keller, G. B. Opposing influence of top-down and bottom-up input on excitatory layer 2/3 neurons in mouse primary visual cortex. Neuron 108, 1194-1206.e5 (2020).

34. Gillon, C. J. et al. Learning from unexpected events in the neocortical microcircuit. 2021.01.15.426915 Preprint at https://doi.org/10.1101/2021.01.15.426915 (2023).

35. Fiser, A. et al. Experience-dependent spatial expectations in mouse visual cortex. Nat. Neurosci. 19, 1658–1664 (2016).

36. Richter, D. & de Lange, F. P. Statistical learning attenuates visual activity only for attended stimuli. eLife 8, e47869 (2019).

37. Richter, D., Ekman, M. & De Lange, F. P. Suppressed sensory response to predictable object stimuli throughout the ventral visual stream. J. Neurosci. 38, 7452–7461 (2018).

38. Blank, H. & Davis, M. H. Prediction errors but not sharpened signals simulate multivoxel fMRI patterns during speech perception. PLoS Biol. 14, 1002577 (2016).

39. Iglesias, S. et al. Hierarchical prediction errors in midbrain and basal forebrain during sensory learning. Neuron 80, 519–530 (2013).

40. Kok, P., Bains, L. J., van Mourik, T., Norris, D. G. & de Lange, F. P. Selective activation of the deep layers of the human primary visual cortex by top-down feedback. Curr. Biol. 26, 371–376 (2016).

41. Press, C., Kok, P. & Yon, D. Learning to perceive and perceiving to learn. Trends Cogn. Sci. 24, 260–261 (2020).

42. Thomas, E. R., Yon, D., De Lange, F. P. & Press, C. Action enhances predicted touch. Psychol. Sci. 33, 48–59 (2022).

43. Press, C. & Yon, D. Perceptual prediction: Rapidly making sense of a noisy world. Curr. Biol. 29, R738–R761 (2019).

44. Yu, A. J. & Dayan, P. Uncertainty, neuromodulation, and attention. Neuron 46, 681–692 (2005).

45. Alilović, J., Timmermans, B., Reteig, L. C., van Gaal, S. & Slagter, H. A. No evidence that predictions and attention modulate the first feedforward sweep of cortical information processing. Cereb. Cortex 29, 2261–2278 (2019).

46. Randeniya, R., Oestreich, L. K. L. & Garrido, M. I. Sensory prediction errors in the continuum of psychosis. Schizophr. Res. 191, 109–122 (2018).

47. Sterzer, P. et al. The predictive coding account of psychosis. Biol. Psychiatry 84, 634–643 (2018).

48. Kafadar, E. et al. Conditioned hallucinations and prior overweighting are state-sensitive markers of hallucination susceptibility. Biol. Psychiatry 92, 772–780 (2022).

49. Haarsma, J., Kok, P. & Browning, M. The promise of layer-specific neuroimaging for testing predictive coding theories of psychosis. Schizophr. Res. 245, 68–76 (2022).

50. Greve, D. N. & Fischl, B. Accurate and robust brain image alignment using boundary-based registration. NeuroImage 48, 63–72 (2009).

51. van Mourik, T., Koopmans, P. J. & Norris, D. G. Improved cortical boundary registration for locally distorted fMRI scans. PLOS ONE 14, e0223440 (2019).

52. Hebart, M. N., Görgen, K. & Haynes, J.-D. The Decoding Toolbox (TDT): a versatile software package for multivariate analyses of functional imaging data. Front. Neuroinformatics 8, (2015).

53. He, H. & Garcia, E. A. Learning from imbalanced data. IEEE Trans. Knowl. Data Eng. 21, 1263–1284 (2009).

