## Supplementary Methods and Results for "Predictions and errors are distinctly represented across V1 layers"

A previous 3T fMRI study of ours, lacking layer-specificity, adopted a similar design to the present study and demonstrated superior decoding of expected relative to unexpected orientations[1]. While it showed no reliable group-level univariate effects, it found that stimulus-preference univariate effects correlated with multivariate effects. We were interested in therefore examining the group-level univariate effects here, alongside their relationship with the multivariate data. The present univariate laminar analysis followed an identical approach to recent studies that separate orientation-specific BOLD responses across layers in V1[2,3] (see Methods).

*ROI definition*

The ROI definition technique was identical to previous studies that determined orientation-specific BOLD activity across layers[2,4–6]. We took the GLM conducted on the localiser task to define the most active V1 voxels and calculated the orientation preference of each of these voxels by contrasting CW and CCW regressors against each other to select the 500 voxels that were most CW preferring (positive *t* values) and CCW preferring (negative *t values*). Separate V1 masks were thus created for CW and CCW preferring voxels. The timecourse of each voxel in these masks was then *z*-scored and multiplied with its *t* value from the orientation contrast, to weight the results towards voxels with the largest orientation preferences[2,4,5].

*Extraction of layer-specific timecourses*

To extract signals from different layers, a spatial GLM approach was adopted to determine the proportion of each voxel's overlap to each of the three grey matter bins in each ROI mask. This approach has previously been used successfully to de-correlate laminar activity profiles in visual areas in tasks of working memory, attention and prediction[2,4,5,7,8], and the present analysis follows an identical method. Described in short, a laminar matrix is designed that represents the distributions of voxels across layers in each ROI and can therefore be used in a spatial GLM to separate BOLD signals for each voxel into the specified layer bins. The GLM of spatially distributed responses unmixes signals from the different layers and produces timecourses for each layer for each ROI in each functional run.

*Estimation of laminar responses*

Timecourses from the three GM layers of interest were then selected for a temporal GLM in SPM12 to estimate laminar responses for each of our conditions. Regressors of interest were modelled for each experimental condition (Expected, Unexpected) and each presented orientation (CW, CCW), resulting in four regressors of interest for each scanning run. These regressors were modelled by convolving stick functions representing the onset of presented stimuli in the main experiment with the canonical haemodynamic response function. Additional nuisance regressors modelled head motion parameters.

To investigate how patterns of activity differed across layers, orientation-specific BOLD responses were calculated for each ROI (e.g., voxels preferring CW in V1), in each run, by subtracting the estimated BOLD activity from conditions that presented the non-preferred orientation (e.g., an expected CCW stimulus) from activity in conditions when the preferred orientation was presented (e.g., an expected CW stimulus). This was done for both expected and unexpected trials, with responses averaged across CW and CCW ROIs, resulting in orientation-specific BOLD responses (preferred activity – non-preferred activity) to each of the conditions in each layer.

### Results

#### *Multivariate and univariate patterns across layers are related*

The ANOVA performed on orientation-specific BOLD responses revealed no significant interaction between Expectation and Layer in V1 ($F(2,40) = .036$, $p = .96$, $np^2 = .002$), alongside no main effects of Expectation ($F(1,20) = 1.77$, $p = .20$, $np^2 = .08$) or Layer ($F(2,40) = .99$, $p = .38$, $np^2 = .05$). The lack of effect at the group level here is perhaps unsurprising given that our 3T study with a similar design also showed no group-level univariate effects. Nevertheless, we imagine that these approaches may be reflecting the operation of broadly similar processes, given our multivariate effects correlated with the univariate effects in our previous work and both analyses reflect orientation-specificity. To examine whether the same was true in this dataset, we examined the correlations between univariate and multivariate effects within each layer. We calculated expectation effect (expected-unexpected) scores for each layer bin for each approach. The Pearson's correlational analysis revealed significant positive correlations between the decoding and univariate approaches for each layer (Deep: $r = .61$, $p = .003$; Middle: $r = .47$, $p = .03$; Superficial: $r = .52$, $p = .016$; see Fig. S1) – reflecting that in each layer participants showing larger expectation effects in decoding show larger univariate stimulus-preference effects, and thus that the two approaches likely examine related information.

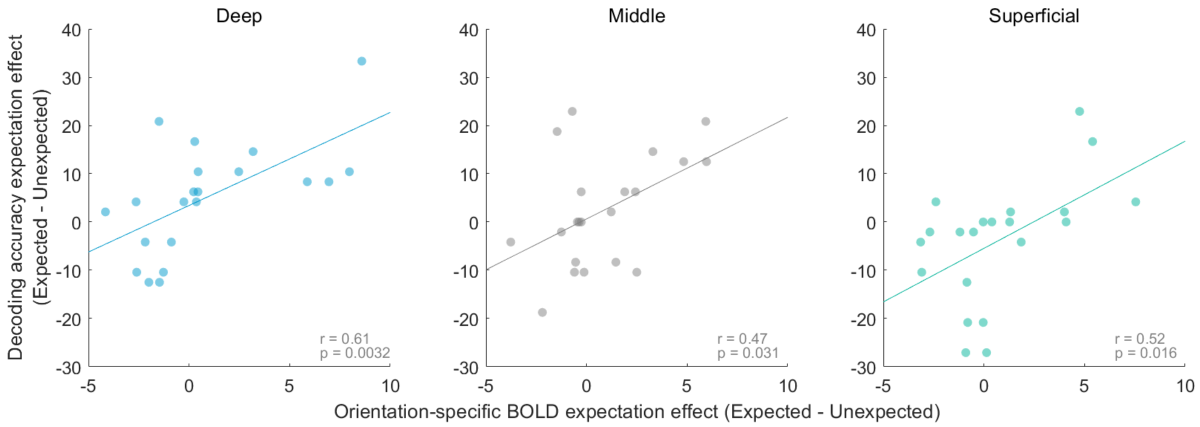


Figure S1. Correlations between the decoding and univariate approaches in each layer bin. Data points represent the expectation effect (expected minus unexpected) for each participant across the two approaches.

### Discussion

Some previous work has used univariate techniques to estimate stimulus-specific activity across layers in visual areas[2–5,8]. While we find in our study that the MVPA analyses are broadly consistent with the univariate method, we do not find significant differences between expected and unexpected events across layers using the univariate method. This suggests that the univariate method might have decreased sensitivity in decoding the representation of stimuli. One explanation is perhaps due to the univariate method using a spatial general linear model to create single time-courses for each layer of interest, thereby controlling for partial volume effects and increasing specificity to layers. However, by reducing the activity of multiple voxels into a single time-course, information about the spatial distribution of individual voxels is lost. In contrast, MVPA uses the information of individual voxels to optimize decoding of expected and unexpected stimuli, but at the cost of having to bin voxels into layers. Despite this MVPA method potentially penalizing on laminar specificity, we were still able to find differential encoding of unexpected events across cortical layers (see Main Text). Future studies could aim to integrate the spatial GLM approach with multivariate methods, to utilize both laminar and stimulus specificity.

### References


1. Yon, D. *et al.* Stubborn predictions in primary visual cortex. *J. Cogn. Neurosci.* **35**, 1133–1143 (2023).
2. Aitken, F. *et al.* Prior expectations evoke stimulus-specific activity in the deep layers of the primary visual cortex. *PLoS Biol.* **18**, e3001023 (2020).
3. Haarsma, J., Deveci, N., Corbin, N., Callaghan, M. F. & Kok, P. Perceptual expectations and false percepts generate stimulus-specific activity in distinct layers of the early visual cortex. *bioRxiv* (2022) doi:10.1101/2022.04.13.488155.
4. Lawrence, S. J. D., Norris, D. G. & de Lange, F. P. Dissociable laminar profiles of concurrent bottom-up and top-down modulation in the human visual cortex. *eLife* **8**, e44422 (2019).
5. Lawrence, S. J. D. *et al.* Laminar organization of working memory signals in human visual cortex. *Curr. Biol.* **28**, 3435-3440.e4 (2018).
6. Albers, A. M., Meindertsma, T., Toni, I. & de Lange, F. P. Decoupling of BOLD amplitude and pattern classification of orientation-selective activity in human visual cortex. *NeuroImage* **180**, 31–40 (2018).
7. Kok, P., Bains, L. J., van Mourik, T., Norris, D. G. & de Lange, F. P. Selective activation of the deep layers of the human primary visual cortex by top-down feedback. *Curr. Biol.* **26**, 371–376 (2016).
8. van Mourik, T., van der Eerden, J. P. J. M., Bazin, P.-L. & Norris, D. G. Laminar signal extraction over extended cortical areas by means of a spatial GLM. *PLOS ONE* **14**, e0212493 (2019).
